# WATLAS: high throughput and real-time tracking of many small birds in the Dutch Wadden Sea

**DOI:** 10.1101/2021.11.08.467683

**Authors:** Allert I. Bijleveld, Frank van Maarseveen, Bas Denissen, Anne Dekinga, Emma Penning, Selin Ersoy, Pratik Gupte, Luc de Monte, Job ten Horn, Roeland A. Bom, Sivan Toledo, Ran Nathan, Christine E. Beardsworth

## Abstract

**Background:** Tracking animal movement is important for understanding how animals interact with their (changing) environment, and crucial for predicting and explaining how animals are affected by anthropogenic effects. The Wadden Sea is a UNESCO World Heritage Site and a region of global importance for millions of small shorebirds. Due to climate change and anthropogenic activity, understanding and predicting movement and space-use in areas like the Wadden Sea is increasingly important. Monitoring and predicting animal movement, however, requires high-resolution tracking of many individuals. While high-resolution tracking has been made possible through GPS, trade-offs between tag weight and battery life limit its use to larger species.

**Methods:** Here, we introduce WATLAS (the Wadden Sea deployment of the ATLAS tracking system) capable of monitoring the movements of hundreds of (small) birds simultaneously in the Dutch Wadden Sea. WATLAS employs an array of receiver stations that can detect and localise small, low-cost tags at fine spatial (meters) and temporal resolution (seconds). From 2017-2021, we tracked red knots, sanderlings, bar-tailed godwits, and common terns. We use parts of these data to give four examples on its performance and how WATLAS can be used to study numerous aspects of animal behaviour, such as, space-use (both intra- and inter-specific), among-individual variation, and social networks.

**Results:** After describing the WATLAS system, we first illustrate space-use of red knots across the study area and how the tidal environment affects their movement. Secondly, we show large among-individual differences in distances travelled per day, and thirdly illustrate how high-throughput WATLAS data allows calculating a proximity-based social network. Finally, we demonstrate that using WATLAS to monitor multiple species can reveal differential space use. For example, despite sanderlings and red knots roosting together, they foraged in different areas of the mudflats.

**Conclusions:** The high-resolution tracking data collected by WATLAS offers many possibilities for research into the drivers of bird movement in the Wadden Sea. WATLAS could provide a tool for impact assessment, and thus aid nature conservation and management of the globally important Wadden Sea ecosystem.

## Background

Movement is a fundamental aspect of life and tracking wild animals under natural conditions has become central to animal behaviour, ecology, and conservation science [1-5]. Animal tracking has revealed extreme and large-scale migratory journeys [6, 7] and detailed patterns of habitat use [8, 9], as well as elucidated mechanisms of navigation [10-12], predator-prey dynamics [13], and social interactions [14]. Insights from animal tracking studies are regularly incorporated in policy and conservation management [5, 15]. For example, identifying important areas for the protection of migration routes [16, 17], detecting wildlife crime [18, 19], and quantifying the human-wildlife conflict [20].

The introduction of the ‘movement ecology’ framework [2], coupled with the rapid development of new tracking technologies and data-processing tools [21, 22] has led to an exponential increase in animal movement ecology research [23]. These developments, particularly the miniaturization of tags capable of generating high-throughput localization data for many individuals simultaneously, allow for novel opportunities to address contemporary questions on individual, group, population, and community level behaviours in the wild [24, 25]. For instance, studies on intra-specific variability [26-28], collective behaviour [29], and interactions among individuals and species with their physical, biotic, and anthropogenic environments [30]. Furthermore, the ongoing miniaturization of tags allows the tracking of ever smaller species, and thus may give a more complete pictures how different species use their habitat [1, 25].

The most common tracking methods, which allow monitoring the movement of animals at high temporal and spatial resolution in the wild, are based on Global Navigation Satellite Systems (GNSS), such as the Global Positioning System (GPS). GPS employs a network of orbiting satellites with known locations, which transmit signals to a receiving tag that uses this information to estimate its position. The principal drawback of GPS is that it does not directly provide a means of reporting position information back to the researcher. The position information is either stored and retrieved later or downloaded via an auxiliary radio-frequency link. In many cases, loggers cannot be retrieved, and the energy costs for transmitting localization data from the animal to the researcher are large [31]. Therefore, trade-offs between data retrieval, sampling frequency, battery size and tag weight often limit the use of these tags and the biological insights gained to larger species [1, 32].

When retrieving loggers is impractical, time-of-arrival (reverse-GPS) and directionof-arrival systems provide an alternative tracking method in which the locations of transmitters need to be estimated and the location of receivers are known. Examples of reverse-GPS systems are presented by MacCurdy et al. [33], Weller-Weiser et al. [34] and Krüger et al. [35]. MOTUS [36] and ARTS [37] are examples of direction-of-arrival transmitter localization. ARGOS [38] is a Doppler-based transmitter-localization satellite system. Because energy costs of transmissions are low and data processing is handled outside of the tag at the receiver end, tags can be small and lightweight while maintaining highthroughput signal transmissions. The smallest VHF-tag compatible with MOTUS, for instance, weighs 0.13 g (www.lotek.com). The spatial resolution of Doppler and direction-of- arrival systems, however, is often coarse. ARGOS, for example, provides a spatial resolution of 250 - 1500 m [38]. MOTUS and ARTS systems can, by triangulating signal-strength of VHF-tag detections, provide localizations at an accuracy of several hundred meters. However, the detection ranges are limited to hundreds of meters up to a few km, and localization error increases with the distance between the tag and receivers [36, 37, 39, 40]. Successful localizations thus require a dense receiver network that limits the size of study area. Time-of-arrival transmitter localization tends to be more accurate, but similarly requires a dense receiver network.

ATLAS (Advanced Tracking and Localization of Animals in real-life Systems) is a recently developed reverse-GPS system [34, 41] that builds on the time-of-arrival wildlife tracking system of MacCurdy et al. [33], and allows real-time tracking and data collection of many small individuals simultaneously. ATLAS comprises an array of stationary receivers that continuously listen for transmissions from small tags. Locations are calculated based on differences in tag-signal arrival times at minimally three receiver stations. Tags are lightweight (0.8 g) and relatively inexpensive (25 €), which facilitates tracking small species and hundreds of individuals simultaneously [42]. Location data is available without retrieval of the tag and in real time, which avoids the need to recapture tagged animals for data retrieval, and allows for locating tagged individuals for auxiliary behavioural observations [43] or for confirming mortality [9]. Whereas GNSS systems and ARGOS allow (near) global tracking, reverse-GPS systems like ATLAS produces location data at a more local scale and, for best performance, requires ‘line of sight’ between receiver and tag [44]. In open landscapes, with published detection ranges up to 40 km for ATLAS [11], its spatial scale is limited by the extent and density of receiver stations. ATLAS has been used to track over 50 different species in four countries, including sites of high scientific or conservation value, such as the Hula Valley in Israel [11, 34] and the Dutch Wadden Sea.

The Wadden Sea is recognized as a UNESCO World Heritage Site for providing a rich habitat for marine mammals [45], fish [46], invertebrates [47], birds [48], and especially migratory shorebirds [49]. Shorebirds form an important component of the Wadden Sea ecosystem, which they use for breeding [50], refuelling during migratory journeys [51], and finding food and safety during their non-breeding periods [52-55]. Millions of shorebirds depend heavily on the worms, snails and shellfish that are found on and in the sediments of the mudflats [56]. Perhaps uniquely, over the past decade, the Wadden Sea has been subject to a large scale benthic macrofauna monitoring survey (Synoptic Intertidal BEnthic Survey (SIBES) [57, 58]), which maps food resources for shorebirds [52, 59]. Combining resource mapping with the simultaneous tracking of many birds offers novel opportunities for studies on space use, trophic interactions and collective behaviour in the wild [60]. Many of the shorebird species are declining in numbers [48], and appear particularly susceptible to the effects of habitat destruction, disturbance, overexploitation of resources, and global climate change [61]. Detailed studies of shorebird space use, in conjunction with knowledge of resource landscapes will offer novel ecological insights and, in combination with monitoring anthropogenic activities, will allow quantifying if and how animals are impacted, which may assist in evidence-based conservation efforts in this important region [62]. Moreover, the Wadden Sea with its flat and open landscape, large numbers of birds, and conservation value, is an ideal candidate for an ATLAS system.

Here, we introduce WATLAS, which is the Wadden Sea deployment of the ATLAS tracking system. In 2017, WATLAS started with 5 receivers and has since grown to have 26 receivers in 2021 (Supplementary Fig. S1 in Additional File 1), making it the largest ATLAS system in the world. The 26 receiver stations are located in the western Dutch Wadden Sea and encompass 1,326 km^2^ (Fig. 1) with a focus on the mudflats surrounding Griend, an important shorebird high-tide roosting site and nature reserve. WATLAS allows simultaneous tracking of several hundred animals at fine temporal and spatial resolution comparable to GPS tracking within a specific study region [63]. So far, WATLAS has been used to track red knots *Calidris canutus* (∽120 g), sanderlings *Calidris alba* (∽50 g), bar-tailed godwits *Limosa lapponica* (∽240 g), and common terns *Sterna hirundo* (∽130 g), but there is scope to track an even larger range of species. In this paper, we will introduce WATLAS, along with several use-cases for the system demonstrating the strengths of the system for studies of movement across levels of organization: from individuals, to species, to populations, and even communities. First, to investigate space use and environmental drivers of movement, we show how space use of red knots tracked in 2019 varies across the entire study area and on a smaller spatial scale across tidal cycles. Second, we give an example of among-individual variation in distance travelled for red knots tagged in 2020. Third, we show how fine-scale movement data WATLAS provides, and how this allows estimating social interactions (proximity-based networks) in red knots. Fourth, as an example of community tracking, we show differences in home ranges between sanderlings and red knots, tracked simultaneously near Richel and Griend. We end by discussing how WATLAS offers possibilities for both fundamental and applied research into the natural and anthropogenic drivers of bird movement in the Wadden Sea.

**Fig. 1.**
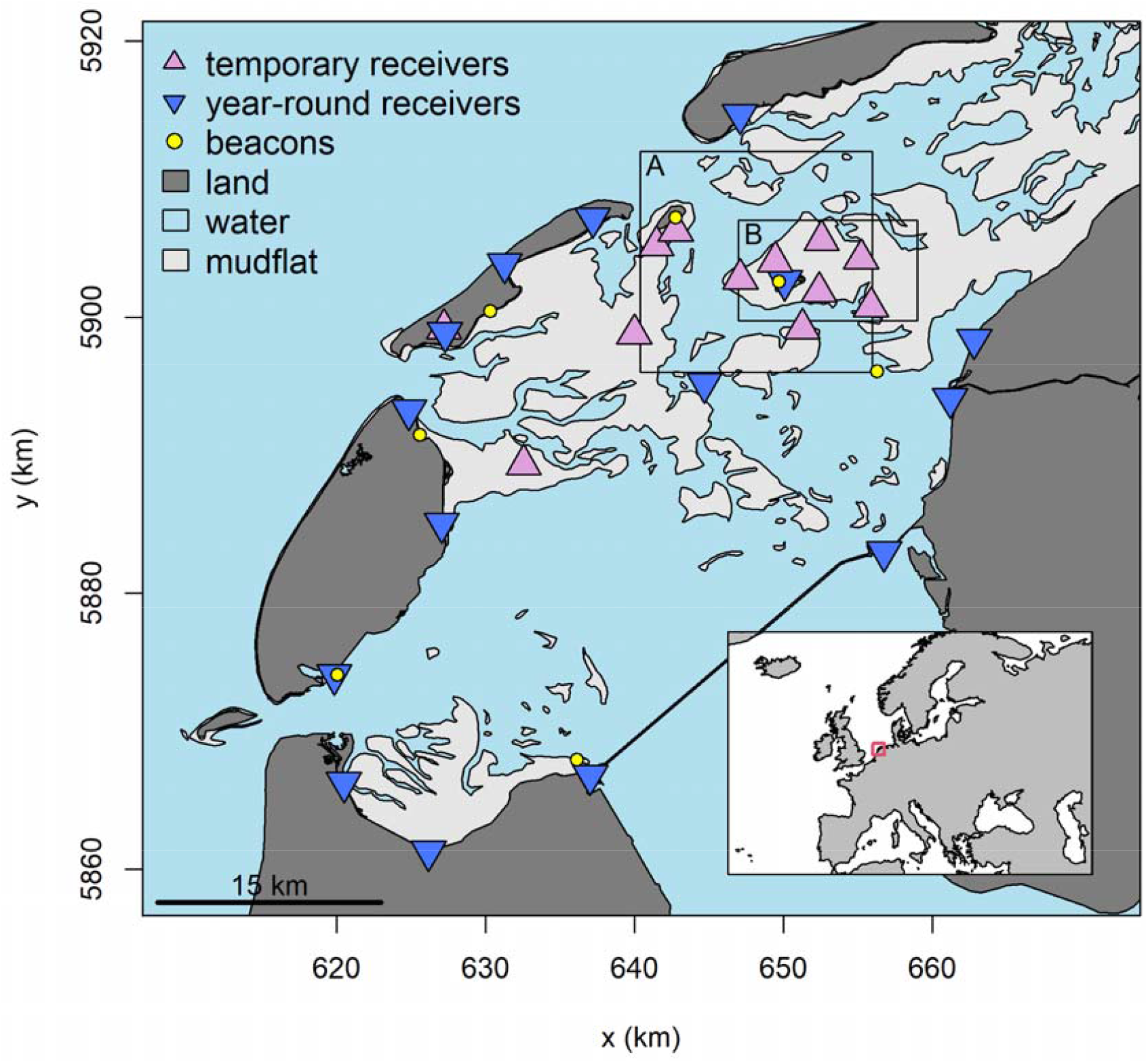
Map of the study area with the receiver array and beacons in 2019. The locations of temporary receivers vary between years (Supplementary Fig. S1 in Additional File 1). Land is shown in dark grey, water in blue and mudflat in light grey. The red square in the inset shows the study area within Europe. Rectangle A shows the area of Richel and Griend as shown in Figures 7 and 9. Rectangle B shows the area of Griend as shown in Figures 4 and 6. The coordinate system refers to UTM 31N. © map data from Rijkswaterstaat.

## Methods

The deployment of WATLAS, which is the Wadden Sea ATLAS system, comprises an array of receivers that continuously listen for tag transmissions. When a transmission is detected, the receiver records the arrival time. These arrival time measurements are sent to a centralized server where location estimates can be computed when at least three receivers detect the signal. Receivers can detect a transmission from any tag in the system at any time, so the tags can transmit as frequently as a localization is needed. Beacons (tags that transmit in fixed known locations – see BEACONS) enable clock-synchronisation across receiver stations.

### RECEIVERS

The WATLAS system currently consists of 26 receiver stations located in the western Wadden Sea (Fig. 1). Fourteen receivers were installed on buildings and other stable structures where power was available, which allowed receivers to be operational year-round. One year-round receiver was placed high on a dune and powered with twelve 100 W solar panels (EnjoySolar) that connected to four 100 Ah AGM batteries (Beaut). Eleven receivers were placed temporarily on the mudflats. Because of the increased likelihood of weather damage in winter, the temporary receivers (Fig. 2) were only in place between July and November each year. One of these temporary receivers was placed on an anchored pontoon that housed a solar powered field station (Fig. 2C). The other ten temporary receivers were attached to scaffolds (Fig. 2A and B) and powered with four 100 W monocrystalline solar panels (EnjoySolar) and a 100 W wind turbine (Ampair), which were connected to three 100 Ah AGM batteries (Beaut). For visibility and safety, a solar powered LED-light was placed on top of the scaffold.

**Fig. 2.**
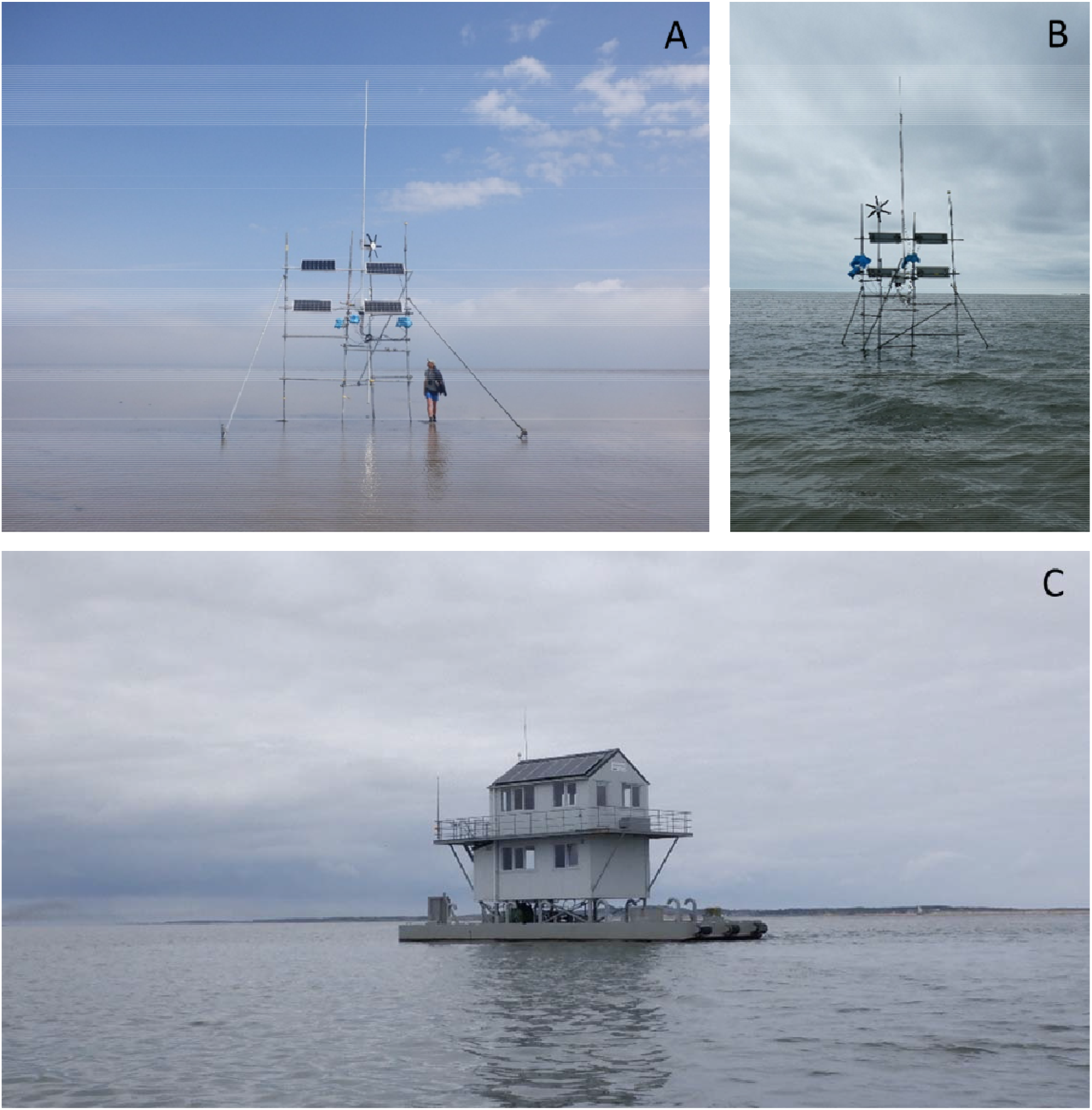
Examples of the temporary solar- and wind powered receiver stations placed on mudflats shown at A) low tide and B) high tide, and C) on the field station.

Each receiver had a 1.5 m Ultra High Frequency (UHF) antenna (Diamond X-50N) mounted on a 6 m aluminium scaffold. To increase the range of tag detections, receiver antennas were placed as high as possible. Antenna height for temporary receivers in 2019 was on average 9.5 m (range: 8.4 – 11.7 m) above sea level, and on average 18.7 m (range: 10.9 to 44.4 m) for year-round receivers.

A coaxial cable connected the antenna to a water-proof cabinet (53 × 43 × 20 cm, Supplementary Fig. S2 in in Additional File 1) via a custom built external Low Noise Amplifier (LNA). The LNA includes a helical bandpass filter to protect against static discharge from thunderclouds. These LNAs connect to a custom front-end unit that acts as a bandpass filter, radio frequency limiter, and power supply to the LNAs [64]. Next comes the Software Defined Radio (SDR) consisting of an USRP N200 with WBX40 daughter board (Ettus Research). This SDR precisely timestamps incoming signal detections using a GPS disciplined oscillator (GPSDO, Ettus Research). The GPSDO was connected to an external amplified ceramic patch antenna (Ettus Research), which allowed the clock rates of all receivers to be synchronized with the atomic clocks from GPS-satellites. Signal transmissions are processed by an onboard computer (Intel NUC i7) that runs Linux. Some computers ran Ubuntu 16 others Ubuntu 20. The number of unique an ATLAS system can detect depends on the processing capability of this Intel NUC. With our onboard computers, we estimate that we can reliably track 300 unique tags that send a transmission every second simultaneously. All receiver stations were connected to the internet using a cellular modem (Huawei E3372 4G/LTE dongle) and an externally mounted antenna (GTT OS-UMTS-0103-C0) to send detection reports to a central server at NIOZ Royal Netherlands Institute for Sea Research. This server runs software that estimates tag locations from time-stamped tag and beacon detections [34]. All data are stored in an online database running MySQL (v5.7, https://www.mysql.com/). Localizations are visualised on www.nioz.nl/watlas in real-time. Moreover, system health (e.g. battery voltage, power usage, and temperature) can also be monitored remotely and in real time for each receiver.

### TAGS

Tags consist of an assembled Printed Circuit Board (PCB), a battery, an antenna, and a protective coating (Fig. 3). The PCBs are based on a CC1310 or CC1350 microcontroller with a built-in Radio-Frequency (RF) transceiver that can transmit a code unique to the tag at 433 MHz. In this study, we used 0.6 g PCB-boards (v2.6.1 and 2.6.2f) [42]. The radio signal is emitted through a 17 cm long antenna made of gold plated multistranded steel wire with a plastic coating, which can handle mechanical stress in a marine environment. Tags were coated with a mixture of two-component epoxy (3M Scotch Weld DP270). To reduce tag weight, the epoxy was mixed with low density glass spheres at a ratio of 1:2. PCB’s are fitted with a (Hall) sensor allowing the tag to be switched on and off with a magnet placed next to the tag. Tags operate at a voltage of 1.8 to 3.8 V and can be fitted with a range of batteries. For example, a pair of silver oxide batteries (0.26 g), single lithium coin-cell batteries ranging from CR1025 (0.7 g) to CR2477 (10.5 g), or a pair of AA batteries (24 g). At signal transmission costs of approximately 0.4 mJ, the capacity of the battery determines the number of transmissions that can be sent, and together with the frequency of transmissions, sets the tag’s operational lifetime (longevity). In 2017 and 2018 we used tag transmission intervals of 1 or 3 s, and in later years 6 s. With WATLAS, we have used CR1620 (1.3 g) and CR2032 batteries (3.0 g), which resulted in final tag weights of respectively 2.4 and 4.4 g (Fig. 3). With a signal transmission interval of 6 s this corresponds to an estimated longevity of 3 and 8 months for the CR1620- and CR2032-loaded tags, respectively (Supplementary Fig. S3 in Additional File 1).

**Fig. 3.**
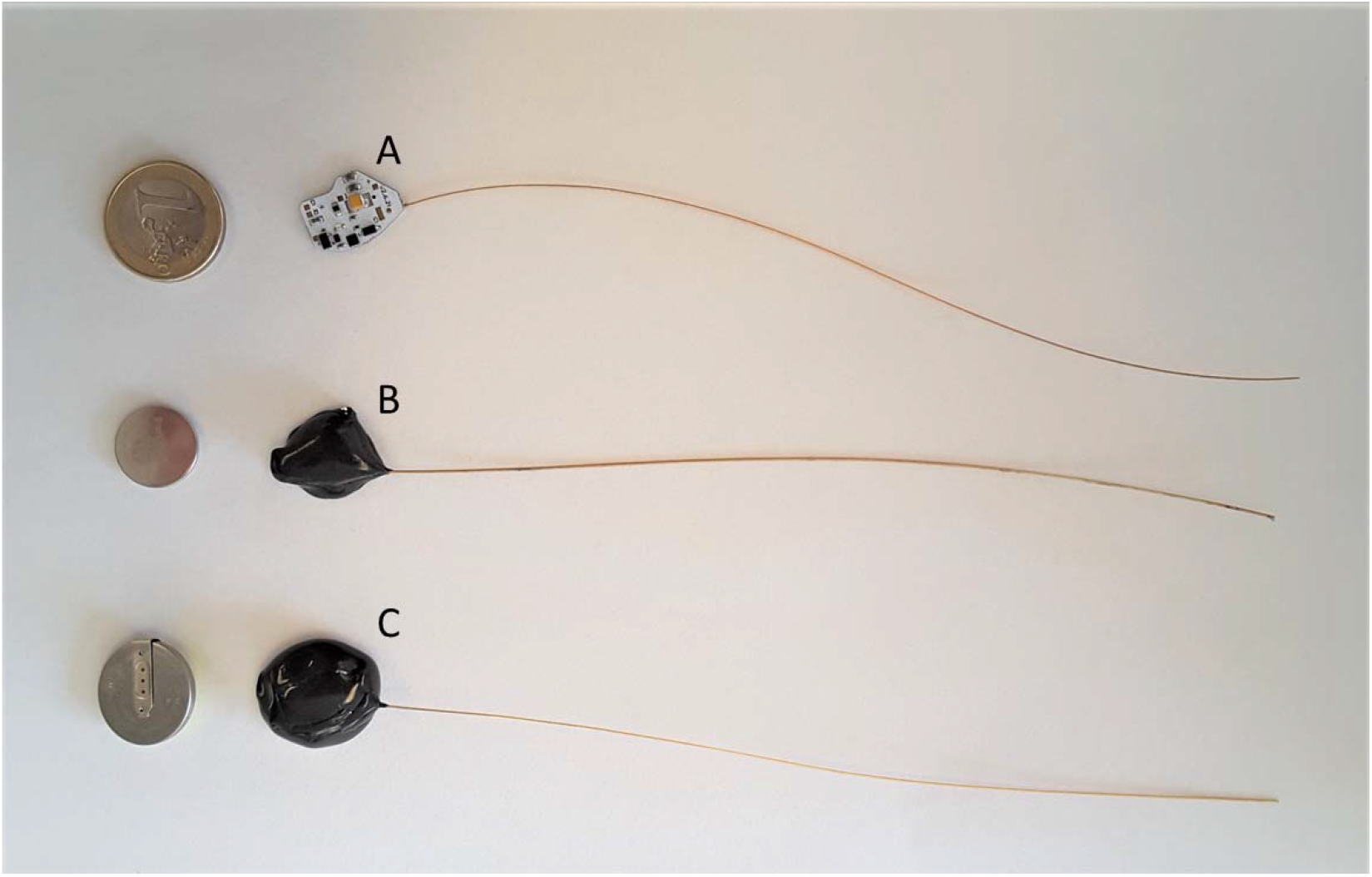
WATLAS tags and batteries. A) tag without battery and coating and a one-euro coin for scale. B) 2.4 g coated tag with CR1620 battery. C) a 4.4 g coated tag with CR2032 battery. The batteries of the tags are shown on the left.

### BEACONS

Beacons were built as standard WATLAS tags, but fitted with one lithium C cell and a helical bandpass filter to protect against static discharge from thunderclouds and connected to a vertical colinear antenna providing 7dB gain in the horizontal plane (Diamond X-50N) identical to the antennas on receivers. The transmission interval of beacons was set to 1 s. Seven beacons were mounted on 6 m aluminium scaffold poles (Supplementary Fig. S4 in Additional File 1). To ensure that each receiver detected at least one beacon consistently, beacons were placed across the study area (Fig. 1 and Supplementary Fig. S1 in Additional File 1). During deployment, the locations of receivers and beacons were recorded with dGPS at 1.5 cm accuracy (Topcon HiPer SR).

### WATLAS COSTS

The most substantial costs are setting up an initial array of receivers. The costs of ATLAS components fluctuate, and more economical configurations are being developed. However, at the time of writing the receiver cabinet with the radio frequency electronics costs about 4,500 €. For a temporary receiver station that requires an independent power supply, there is an additional cost of 5,000 € that includes equipment for generating wind and solar power, batteries, and scaffolding. Tag cost is dominated by costs of assembling the electronics, and this largely depends on the numbers of tags produced in a batch: 100 € each at 20 pcs and 22 € each at 200 pcs (www.circuithub.com). The labour costs of tag assembly can easily cost an equal amount. Operational costs can be quite substantial as well, such as those for mobile data transfer. For example, between August and November 2018 receivers transferred an average of 14 GB of data per month (7 to 18 GB per receiver per month). Per receiver, the monthly costs for an unlimited data plan were 35 €.

### TRACKING SHOREBIRDS WITH WATLAS

We present examples from red knots tracked in 2018 (N = 79), 2019 (N = 221) and 2020 (N=44), and sanderlings tracked in 2018 (N = 35). Red knots were caught on Richel (53.28° N, 5.01° E) and Griend (53.25° N, 5.25° E) (Fig. 1) with mist nets during new moon periods each year between July and October. Most sanderling (N = 33) were caught on Griend by means of canon netting on 26 July 2018, but some by mist-netting on 12 August 2018 (N = 2). All birds were banded with unique combinations of colour-rings and released after gluing a WATLAS tag to their rump with cyanoacrylate glue (Fig. 4). Red knots were fitted with 4.4 g tags (Fig. 3C) that were on average 3.2 % (SD = 0.2) of body mass. Sanderling were fitted with 2.4 g tags (Fig. 3B) that were on average 4.4 % (SD = 0.4) of body mass. Because red knots and sanderlings can double in body mass while fuelling for migration, we believe the additional weight of the tag (<5% of body mass) did not cause substantial detrimental effects [65, 66]. All birds were released from Richel and Griend.

**Fig. 4.**
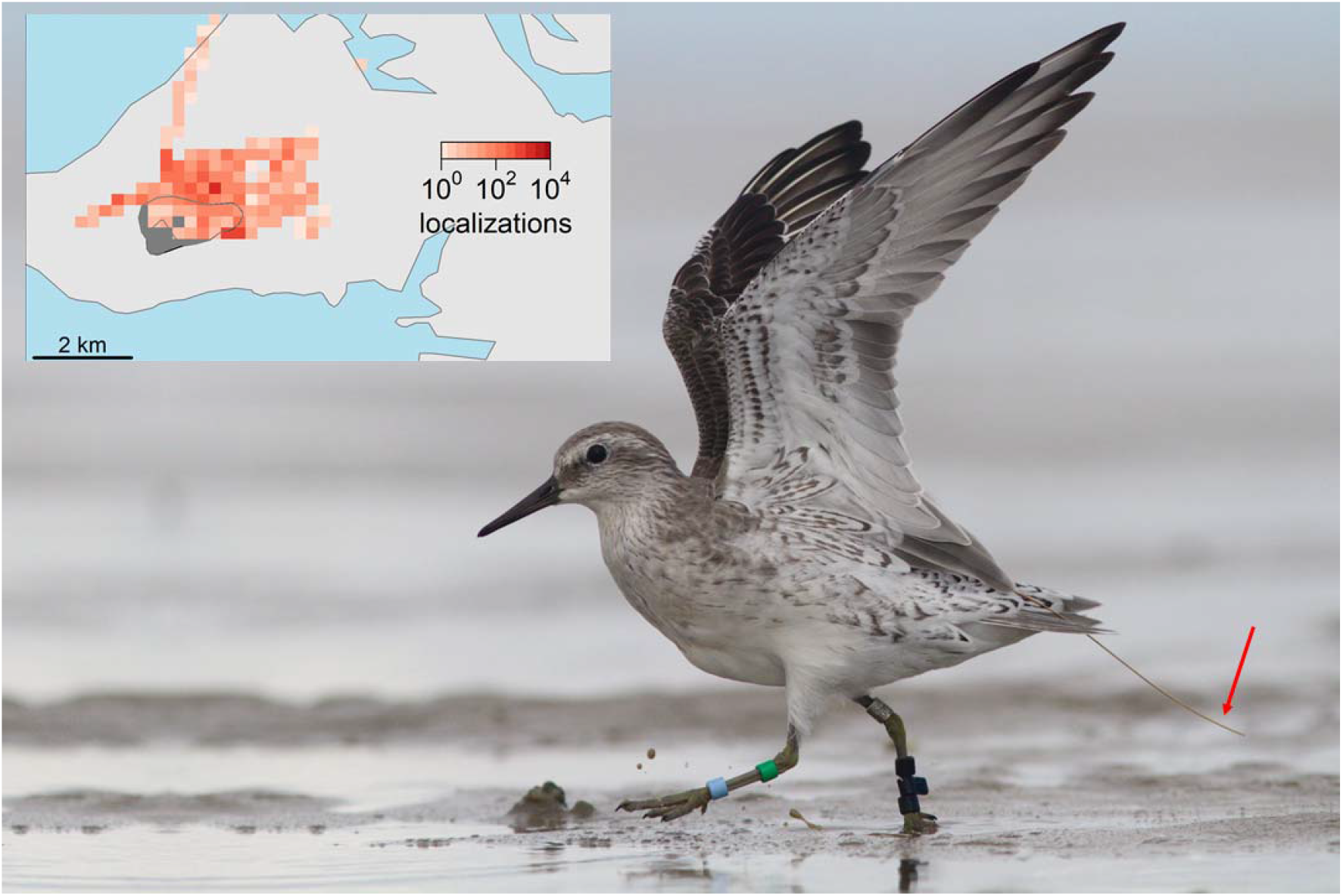
A colour-ringed red knot in winter plumage, bearing a WATLAS tag glued to its rump; the tag antenna can be seen extending beyond the tail to the right of the image as indicated with the red arrow. WATLAS tags allow free movement of the wings and fall off as the feathers underneath the tag regrow. The inset shows this bird’s localizations around Griend, collected between 15 September and 21 September 2017. See rectangle B in Fig. 1 for placement of the inset within the study area. © map data from Rijkswaterstaat, and photo taken on 16 September 2017 by Benjamin Gnep.

### ACCURACY OF WATLAS AND TRACKING DURATION

In another study [63], the bias, accuracy and precision of WATLAS was quantified. In brief, WATLAS location estimates were comparable to GPS with localization errors of 4 - 6 m. Accuracy was higher if more receivers detected the tag and localization error was reduced to a few meters. The tag’s location in respect to the receiver array configuration had a large effect, with less accurate estimates occurring when the tag was on the outskirts or outside the array of receivers. The proportion of localizations of a stationary test tag 1.2 m above ground was over 90% of the expected fixes, but for tagged red knots on the ground this proportion reduced to 51%. Additionally, the proportion of accurate localizations of tagged birds was higher when they were central to the array, and lower on the outskirts of the array [Fig. 6 in 63]. Nonetheless, with the small interval between tag transmissions, a fix rate of 51% still resulted in near continuous tracking, and even near the edges of the array provided reliable and useful localizations.

At a 6s interval, the theoretical longevity of the 4.4 g tags attached to red knots was 8 months. Indeed, this matched the maximum duration of 239 days that red knots were detected in 2019 (Fig. 9 in [42]). Fig. 9 in Toledo et al. [42], however, also shows a steady decline in the number of unique tags tracked, which was particularly pronounced directly after release. Reasons for this decline are multiple and include tags failing, birds dying, and birds leaving the tracking area. Another important aspect is that the period of tagging coincided with the period that red knots moult into their winter plumage [67]. Because we glue the tags to the birds, birds in moult will lose their tags and localizations will cease. In 2021, we recaptured a red knot that was tagged the previous year and did not show visible signs where the tag had been attached.

### PRE-PROCESSING WATLAS DATA TO IMPROVE POSITION ESTIMATES

The accuracy of WATLAS localizations is comparable to conventional GNSS systems [63] and to the Hula Valley ATLAS system [34]. However, in common with other positioning systems, WATLAS data can contain some inaccurate localization estimates. Sources of the localization errors can be due to temporary mismatches in clock synchronization between receivers, RF-interference, or tag signal collisions. To reduce errors in positioning estimates, filtering and smoothing localization data is common practice in movement ecology [68]. Here, we used a simple filter-smoothing process on the estimates that are reported by the localization algorithm, namely variance in the Easting and Northing. We removed localizations that had variances in Easting and Northing above 2,000 m^2^. Additionally, we smoothed the data with ‘runmed’ in R [69] by computing a 5-point median smooth across the localizations [52, 68]. After data processing the precision and error of localizations can be reduced to several metres [63].

## Results

We will demonstrate examples for how WATLAS opens-up possibilities for studying space-use and environmental drivers of movement, among-individual variation in distance travelled, intra-specific (social) interactions, and community tracking with inter-specific space use in the wild. All analyses were done in R v4.0.2 [69].

### EXAMPLE 1. ESTIMATING SPACE-USE

To show how WATLAS can be used to investigate space use and e.g. identify hotspots, we created heatmaps of the localisations of 221 red knots tracked between 1 August 2019 and 1 November 2019 (92 days). We created heatmaps at two spatial scales: The large spatial scale of the entire study area in grid cells of 500 × 500 m, as well as the smaller spatial scale around Richel and Griend with grid cells of 250 × 250 m. To additionally illustrate how WATLAS data can be used to investigate environmental drivers of space use, we created heatmaps on a smaller spatial scale separately for the different phases of the tidal cycle. The tidal-phases were selected based on the water level (NAP; Amsterdam Ordnance Datum) at the tide gauge at West-Terschelling including a delay of 30 min due to the distance from Griend (53.37° N, 5. 22° E): high tide (> 100 cm NAP), first ebb tide (outgoing tide between 50 and 100 cm NAP), second ebb tide (outgoing tide between -50 and 50 cm NAP), low tide (< -50 cm NAP), first flood tide (incoming tide between -50 and 50 cm NAP), second flood tide (incoming tide between 50 and 100 cm NAP). These tags were programmed to transmit every 6 s, thus each location represents at least 6 s of space use.

The large-scale heatmap confirmed that Richel and Griend, where the knots were caught, are hotspots. Nonetheless, red knots spread-out across the entire study area (Fig. 5). It should be noted, however, that the number of localizations, being related to the density of receiving stations, is higher in the central part of the area than on the border. As can be seen from the localizations over relatively deep gullies, several birds moved between islands and the mainland (Supplementary Fig. S5 in Additional File 1). Likewise, localizations across the North Sea suggest that birds crossed in the direction of the United Kingdom. In some cases, these birds were detected up to 34 km from the closest receiver.

**Fig. 5.**
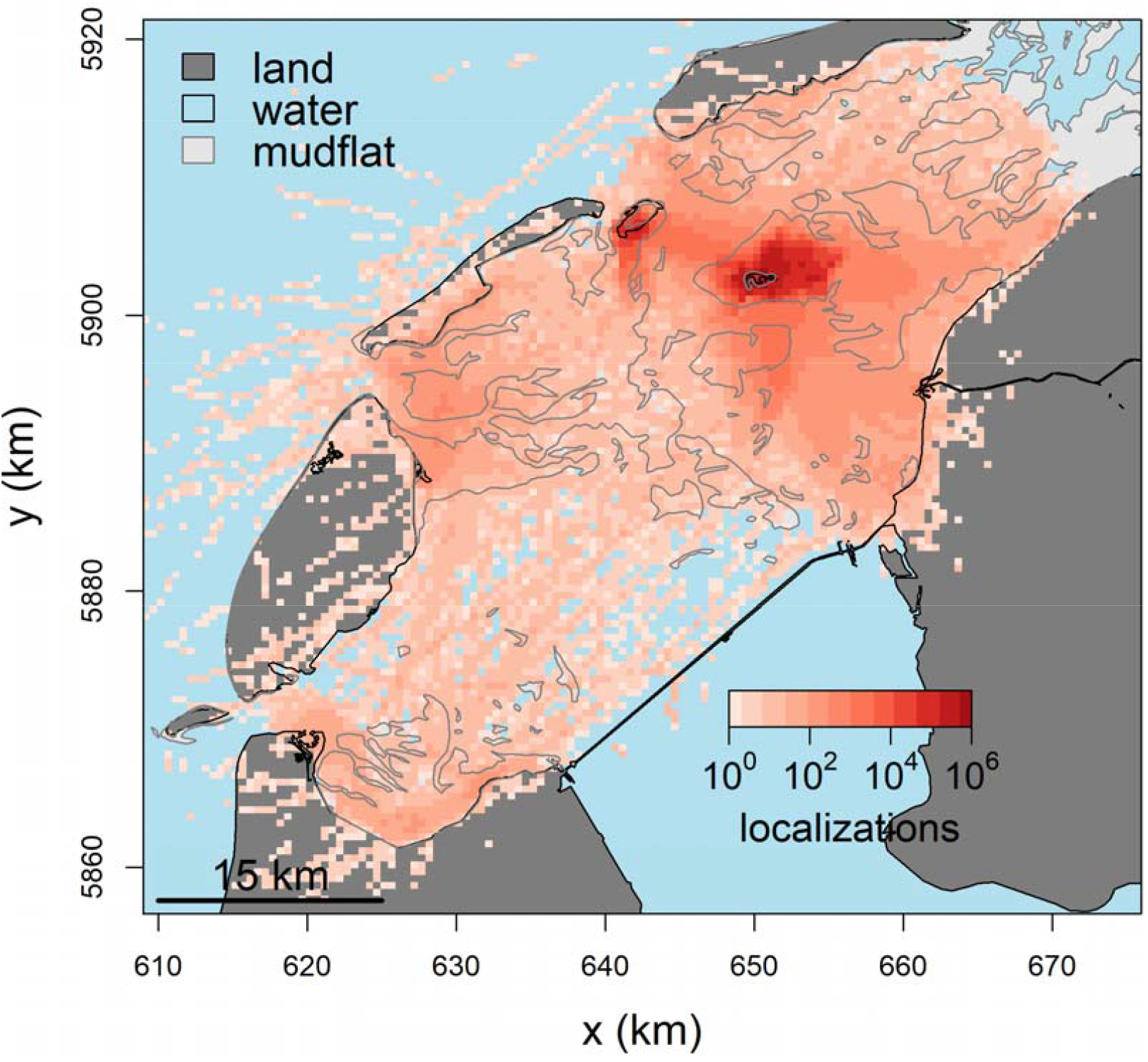
Large-scale space use of 221 red knots tracked between 1 August and 1 November 2019 (92 days) within the entire study area. The colour scale represents the number of localizations in 500 × 500 m grid cells. Note that the colour scale is logarithmic. Water is coloured blue, land dark grey, and mudflats light grey with a solid line indicating their boundery. Because the tags send a signal at 1/6 Hz, each localization represents a minimum of 6 s of space use for red knots. The coordinate system refers to UTM 31N. © map data from Rijkswaterstaat.

On a smaller spatial scale, the heatmaps for different phases of the tidal cycle around Richel and Griend showed how the tidal dynamic affects space use of red knots on a population level (Fig. 6). With the outgoing tide, the birds moved out on the now-exposed mudflats in search of invertebrate prey. Interestingly, space-use differed between ebb and flood tides even though the water level was the same. This can, for instance, be seen by comparing Fig. 6C with 6E, and Fig. 6B with 6F. The fewest localizations were observed during the low tide period, probably because birds spread out and even moved outside the tracking area. With the incoming tide the birds returned and aggregated on Richel and Griend.

**Fig. 6.**
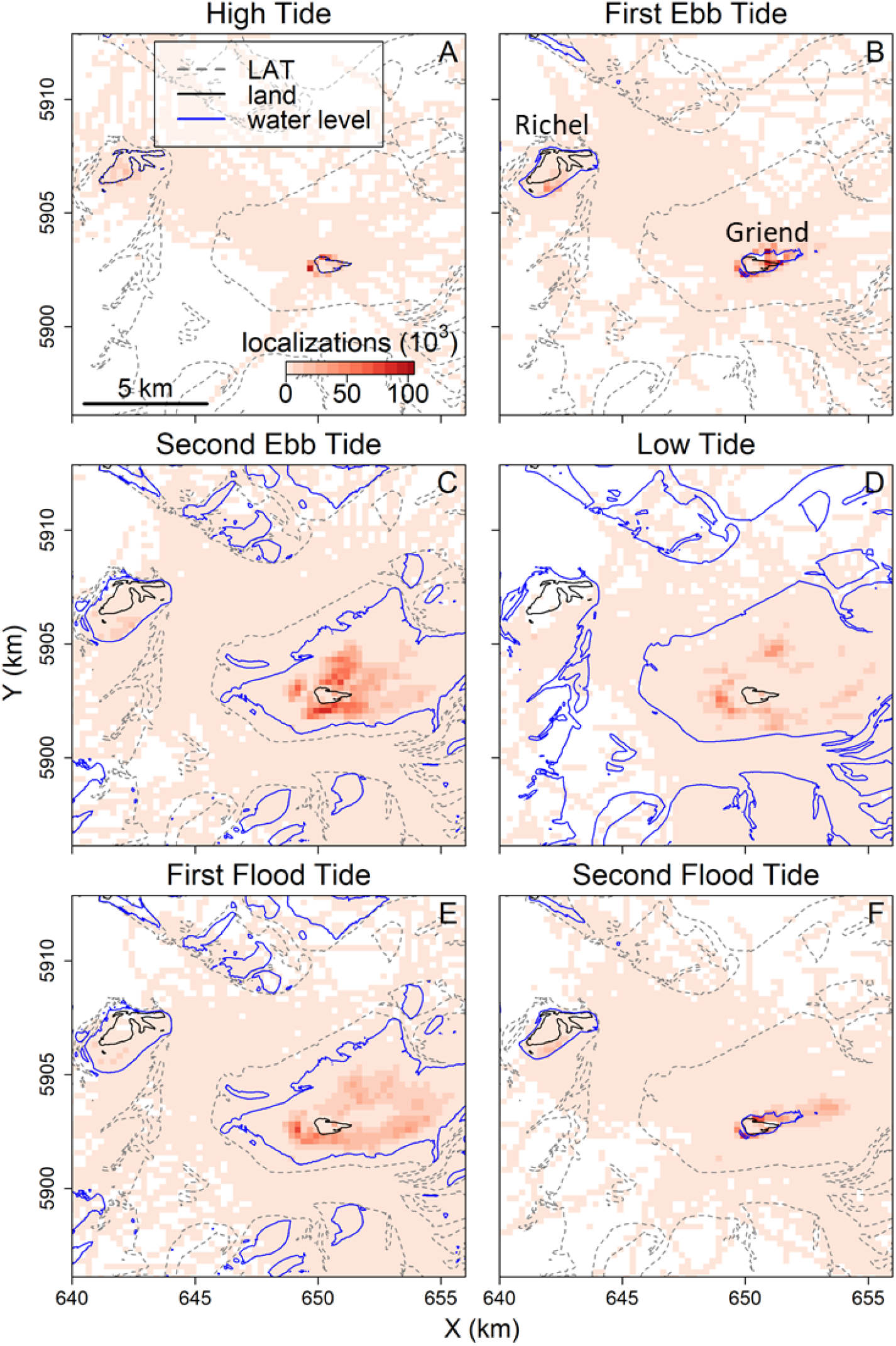
Small-scale space use during different phases of the tidal cycle for 221 red knots near Richel and Griend (locations labelled in panel B) between 1 August and 1 November 2019 (92 days). Panels show different phases of the tidal cycle from high tide (panel A), through ebb tide (panels B and C), to low tide (panel D) and flood tide (panels E and F). The colour scale represents the number of localizations in 250 × 250 m grid cells. The boundary of mudflats are indicated with a grey dashed line (i.e. the Lowest Astronomical Tide, LAT). The blue line indicates the lowest water level within the different tidal phases. Land is indicated with a solid black line. Because the tags send a signal at 1/6 Hz, each localization represents a minimum of 6 s of space use for red knots. See rectangle A in Fig. 1 for placement of this map within the study area. The coordinate system refers to UTM 31N. © map data from Rijkswaterstaat.

### EXAMPLE 2. AMONG-INDIVIDUAL VARIATION IN MOVEMENT

To illustrate the large-scale application of WATLAS tracking and to explore among-individual variation in space use, we selected data from seven red knots (out of 44) on 17 October 2020. These tags were programmed to transmit every 6 s. Additionally, we calculated cumulative distance between successive localizations to reveal variation in daily distances travelled for all individuals tracked that day.

Red knots were successfully localized in large parts of the study area, though gaps in the tracks also occurred (Fig. 7). These gaps happened especially where the density of receivers was low, i.e. when the tag was not within the detection range of at least three receivers. The tracking data revealed substantial differences among individuals in the distance travelled over a 24-hour period, which ranged between 42 and 328 km day^-1^ for all 44 birds (mean±SD: 131±49 km d^-1^, histogram in Fig. 7). There were also differences in the number of localizations between birds (mean = 2,111 bird^-1^; range = 144 - 3,680), which significantly explained distance travelled per day (slope = 26.3 m per localization, p<0.01). Nonetheless, when dividing the distance travelled by the number of localizations per bird, there were still large among-individual differences (mean = 83.1 m per localization; range = 27.3 - 316.5), which shows that the among-individual variation in distance travelled is not merely caused by differences in the number of successful localizations.

**Fig. 7.**
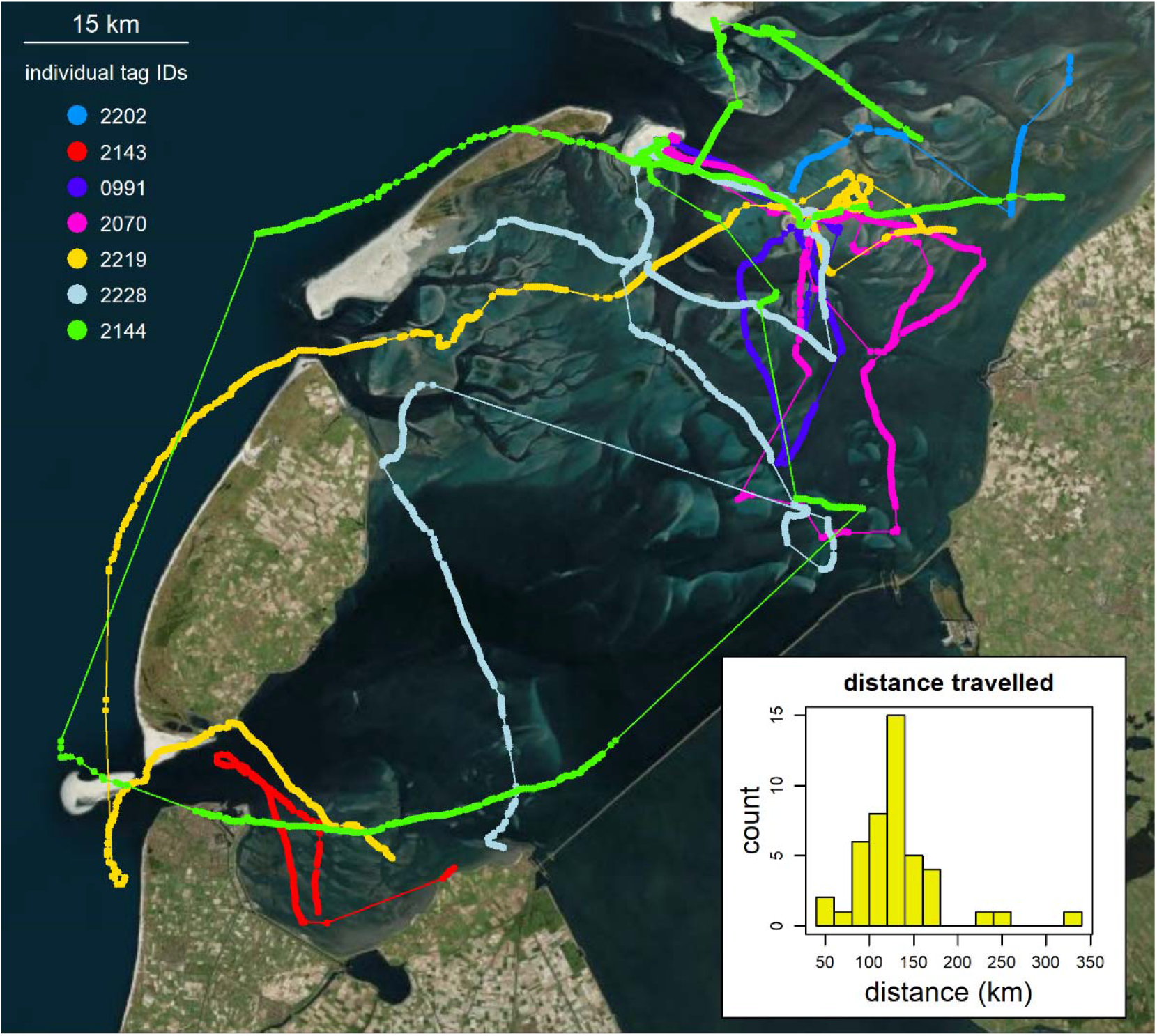
Tracks for a subset of seven individual red knots that differ in the spatial scale of space use across the entire study area. Data collected over 24 hours on 17 October 2020 are shown. The inset shows the histogram of cumulative distance travelled for all 44 birds tracked on this day. © map data from Bing, OpenStreetMap.

### EXAMPLE 3. FINE SCALE MOVEMENT AND INTRA-SPECIFIC INTERACTIONS

To illustrate the application of high-resolution WATLAS data for investigating social interactions, we selected seven red knots (out of 79) tracked near Griend (Fig. 1) between two high tides at 00:50 and 13:16 CEST on 31 August 2018. These data, recorded at 1/3Hz, were aggregated into 30 s timesteps, and the mean coordinates were calculated. Within these time steps, social proximity was defined as being within 50 m of each other [70]. The social network was created in R with the library ‘spatsoc’ [71].

The fine-scale movement patterns of the red knots confirmed (Fig. 8) that birds roosted on Griend during high tide and, as the water recedes with the ebb tide, moved out onto the exposed mudflat to forage. While foraging, the birds walked across the mudflat, which resulted in areas with dense localizations. Birds flew between different areas to forage as can be seen by the areas of dense localizations connected by lines with sparser localizations (flight). An animation of the fine-scale movement with the incoming tide can be found in the supplement (Additional file 2). Tracking the fine-scale movements of many tagged animals allows for the investigation of inter-individual interactions. For instance, the proximity-based social network of our subset of seven red knots revealed that some individuals were often in close proximity (e.g. birds with tag IDs 409 and 412; Fig. 8), whereas some individuals were rarely close to the other individuals. The individual with tag ID 458 was mostly static, hence rarely close to any other tagged individual within this period. The comparison between the tracks and proximity network further shows the merit of collecting fine-scale high-resolution tracking data. Visually, for example, the individual with tag ID 409 seems to have much higher overlap with the individual with tag ID 418 than with the individual with tag ID 412. Nonetheless, the spatiotemporal proximity network shows that individuals with tag IDs 409 and 412 have the highest overlap.

**Fig. 8.**
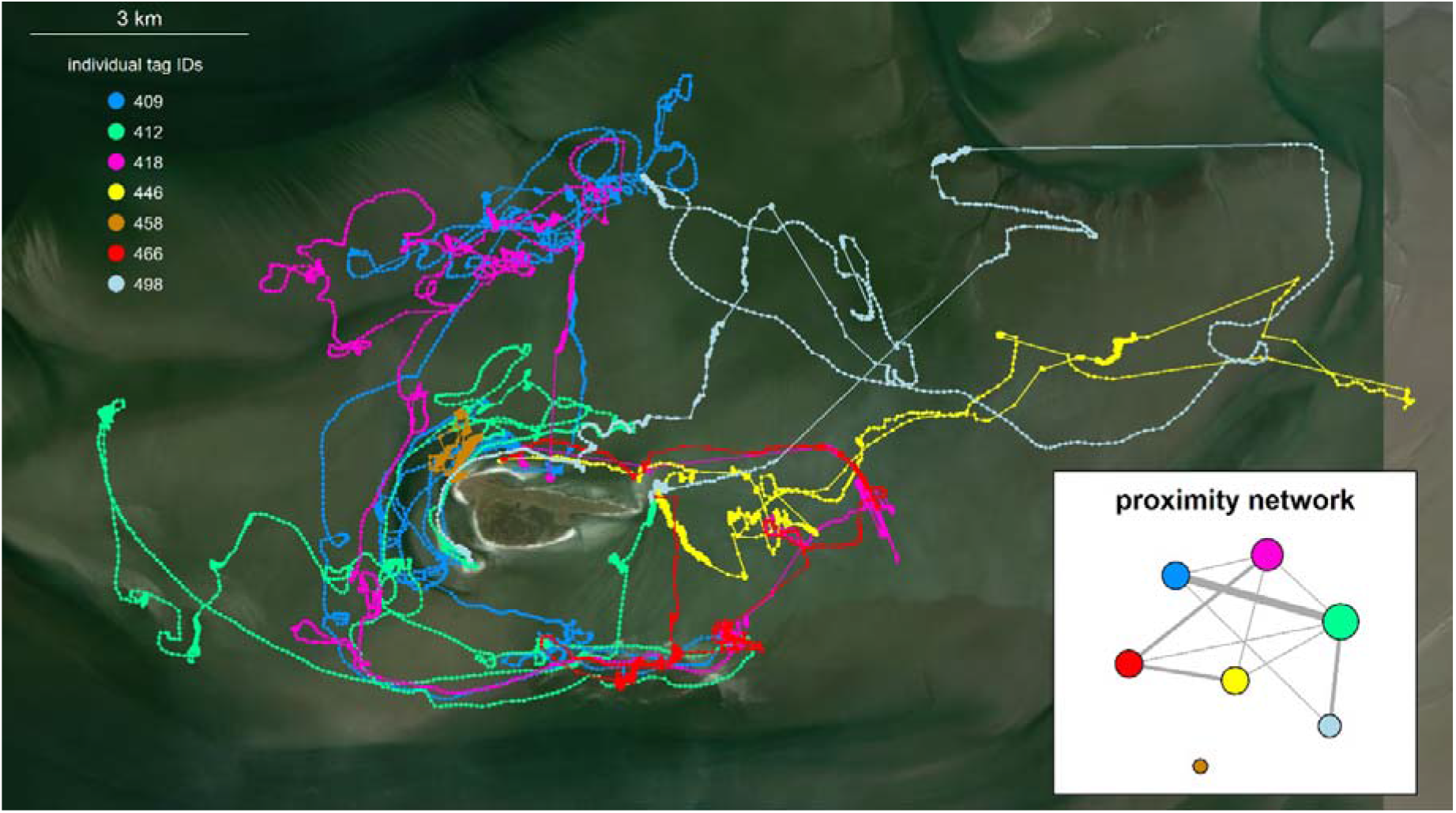
Detailed movements from a subset of seven tracked individual red knots around Griend. Data between two high tides at 00:50 and 13:16 CEST on 31 August 2018 are shown. The inset shows the proximity network for these seven birds based on a spatial proximity of 50 m (see methods). In total, 79 individuals were localized in this timeframe for which an animation of their movement relative to the tide can be found in the Supplementary Information. See rectangle B in Fig. 1 for placement of this map within the study area. © map data from Bing, OpenStreetMap.

### EXAMPLE 4. COMMUNITY TRACKING

Tracking individuals from different species within one region, allows investigations of inter-specific space use. To show how space use, comprised of individual movements, scales up to community-level space use, we analyse differences in the home ranges of sanderlings and red knots. Kernel densities were calculated with the R-library ‘amt’ [72] for 35 sanderling and 35 red knots tracked simultaneously between 12 and 19 August 2018. More red knots had been successfully tracked in that period, but to equalize sample sizes with the sanderlings, were randomly selected. Note that the mean number of localizations in this period for sanderlings was 1,294 fixes (SD 2,745) and that for red knots was 9,521 fixes (SD 9,868).

The home-range analyses show that although sanderlings and red knots both roost on Richel and Griend, they differ in their low-tide distribution (Fig. 9). The home range of sanderlings appeared larger than that of red knots and included intertidal flats near Richel and extended more to the east of Griend. The differences in space use between sanderlings and red knots might be related to differences in the behaviour and spatial distribution of their prey. For instance, red knots forage on patchily distributed and relatively sessile shellfish, whereas sanderlings forage on shrimp that are mobile and follow the tide.

**Fig. 9.**
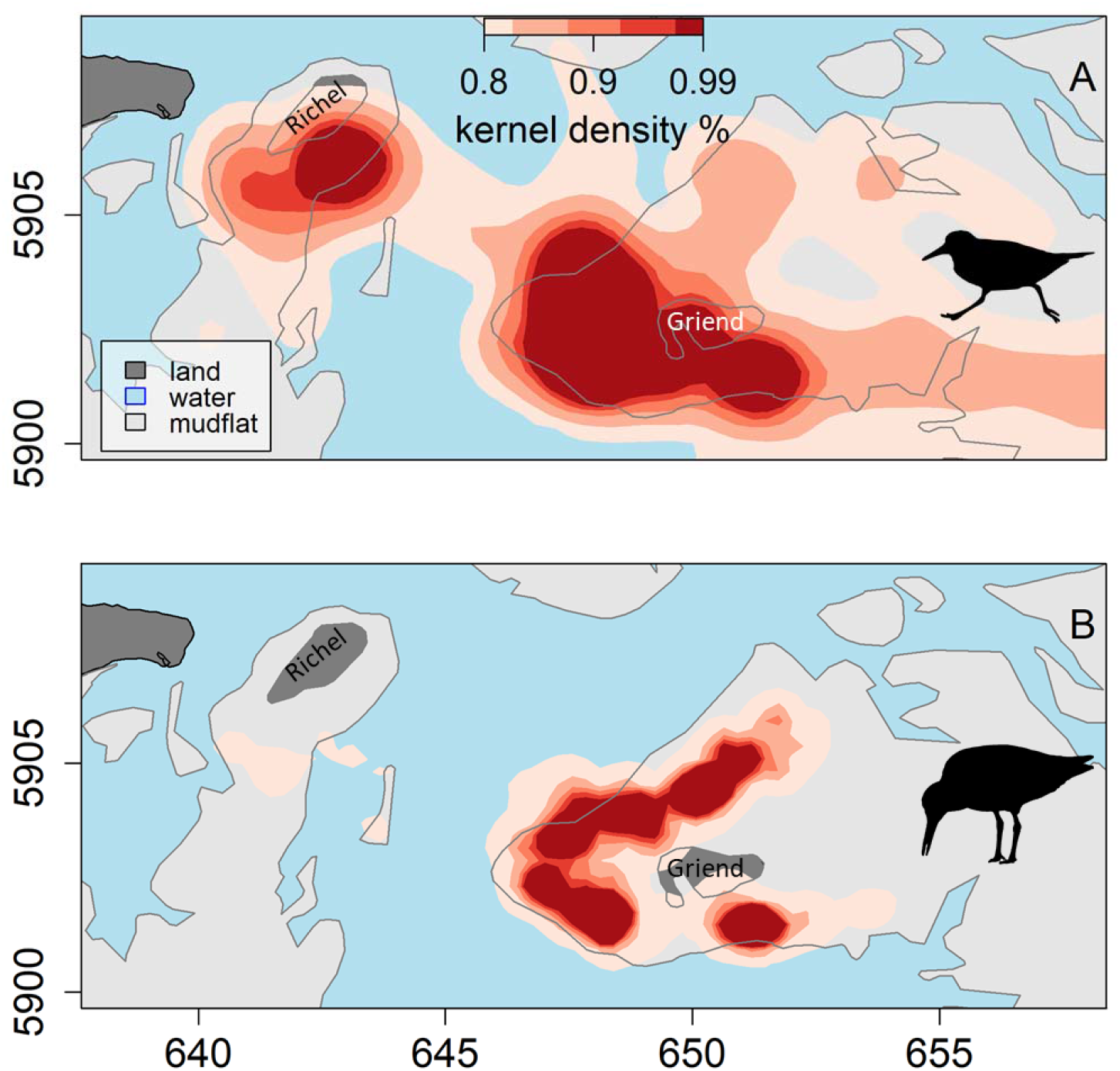
Space use by sanderlings and red knots. The colour scale shows home range estimates with kernel densities for A) 35 sanderlings and B) 35 red knots during low tide between 12 and 19 August 2018. The islets of Richel and Griend are roosting sites for these sanderlings and red knots. Water is coloured blue, land dark grey, and mudflats light grey. See rectangle A in Fig. 1 for placement of this map within the study area. The coordinate system refers to UTM 31N. © map data from Rijkswaterstaat.

## Discussion

With an array of 26 receiver stations located in the western Dutch Wadden Sea, WATLAS covers 1,326 km^2^ and is currently the largest deployment of an ATLAS tracking system worldwide. With examples from red knots and sanderlings, we illustrated various applications of the high spatial and temporal resolution movement data obtained by WATLAS. Moreover, we provided examples of how high-throughput movement data can be utilized to study important aspects of animal movement ecology and space use, such as among-individual variation in behaviour, intra-specific interactions (social networks) as well as inter-specific interactions (community assembly). For regional-scale studies on small animals, reverse-GPS systems like ATLAS, are promising.

### TECHNICAL CONSIDERATIONS

For successful localization, reverse-GPS tracking like ATLAS, requires at least three receivers to detect the tag’s signal, and signal detection requires a ‘line of sight’. A study on the accuracy of WATLAS localizations, showed that tags (∽1.2 m from ground) were mostly detected by receivers within 5 km of the tag [63]. Near Richel and Griend, where the distances between receivers were smallest, localizations were most numerous. Near the edges of the array and on the large-scale of the entire study area, where the distance between receivers was largest (Fig. 1), tags were localized less often, causing gaps in the tracks (Fig. 7). To avoid missing localizations, the density of receivers can be increased, and the array should surround the main area of interest [63].

Tags attached to animals in flight generally have larger detection ranges than animals on the ground, due to their usually greater height. For instance, in another ATLAS system [11], Egyptian Fruit Bats *Rousettus aegyptiacus* were detected during flight up to 40 km away from the receivers. In our study system, we recorded similar detection ranges of birds in flight, and we were able to localise them across the entire study area up to 34 km from the nearest receiver.

The most substantial costs for reverse-GPS are setting up the initial array of receivers. Once a reverse-GPS tracking system is in place, the relatively cheap tags allows tracking large and representative samples of animal populations at low costs [31]. The number of unique tags ATLAS systems can detect simultaneously is limited, but so far WATLAS has tracked 232 tags simultaneously without problems. This limit is set by the processing capability of the computer within the receiver, as well as interference of overlapping transmissions between tags. The percentage of missed tag transmissions increases exponentially as a function of the number of transmitters within range (see Fig. 3.5 in [73]). Note that both limitations are not an intrinsic limitation of ATLAS, but a limitation of the current implementation. More powerful processors in the receivers will, for instance, allow more and simultaneous tag detections.

Another advantage of ATLAS systems is the weight of the trackers. So far, the lightest ATLAS tag (including battery and coating) weighs as little as 0.8 g [42], which allows tracking small and light-weight individuals that were previously too small to track remotely at high spatial accuracy. Other tracking systems that allow tracking of smaller free-living individuals, include MOTUS [36], and light-level geolocation data loggers [74]. These devices can provide high temporal resolution data or be used to track birds over large areas. Compared to ATLAS, however, the spatial accuracy of localizations with MOTUS and geolocation loggers is large (kms) and retrieving data from geolocation loggers requires recapturing the tracked animals. Recapturing can be problematic and prevents real-time observations and analyses of tracked animals. Another promising tracking system is ICARUS [75, 76], but this is under development and tags are estimated to be larger than the lightest tags of ATLAS.

Compared to GPS systems, reverse-GPS systems have dramatically reduced tag energy consumption, which allows high temporal resolution localizations to be collected with small batteries [31] and tags can be expected to last for longer for a given number of localizations [73]. Because of the eight months maximum lifetime of WATLAS-tags used in this study, the tags are glued to the backs of birds and will fall off during body moult. From an ethical perspective this is preferred over e.g. full body harnesses [77], because animals need to cope with the added weight [78] and potential aerodynamic discomfort only temporarily [79].

Despite the promise of ATLAS, it is necessary to note some limitations of the system. Compared to other more global tracking systems, ATLAS is limited to much smaller scales and therefore is unsuitable for some study designs. Furthermore, the ATLAS system was developed and is being applied by multidisciplinary teams of scientists for movement ecology research [41]. Each system has been installed and is maintained by scientists, and often requires collaborations with landowners and organisations for establishing receiver locations. This is in stark contrast to plug-and-play tags like many GPS or ARGOS tags. Other biologgers also collect auxiliary information, such as altitude and acceleration [21]. While not in use (yet) in our WATLAS system, ATLAS tags have also been successfully fitted with air-pressure, temperature, and humidity sensors [42] and an on-board accelerometer is under development. However, remote transfer of data from such sensors remain challenging.

### OPPORTUNITIES FOR ANIMAL MOVEMENT RESEARCH AND CONSERVATION

Simultaneously tracking many free-living small birds at high spatial and temporal resolution, allows for novel studies on e.g., among-individual variation, collective behaviour, and inter-specific interactions in the wild [60]. Moreover, because individuals from different species and different trophic levels can be tracked simultaneously, exciting opportunities exist for studying movement ecology at the community level [80]. Clearly, the list of examples given here is not exhaustive for what is possible with an ATLAS system like WATLAS. Many individual or environmental factors can be linked with movement data which can be used to describe or predict behaviour. Specifically, within the Dutch Wadden Sea, of particular interest is the ability to combine shorebird tracking with knowledge on their food resources. In this study, we illustrated differences in home range between sanderlings and red knots. Because sanderling prefer shrimp and red knot prefer shellfish [55], the differences in space-use are likely related to differences in the distribution of their preferred prey. Linking space-use of these birds with the large-scale mapping of benthic resources in collaboration with the SIBES project [57, 58], will offer exciting opportunities for understanding consumer-resource interactions and space-use in intertidal ecosystems. Having long-term tracking systems like WATLAS established also allows researchers to sample behaviour opportunistically. In 2020, more than 3,000 red knots with a highly pathogenic avian influenza virus were found dead in the Wadden sea [81]. Avian influenza is a disease of global concern, which can affect both domestic and wild populations. However, none of the found red knots were tagged with WATLAS but collecting data on movement and social interactions could provide valuable insights into disease transmission [82-84].

Despite strong legal protection and management measures being in place, many activities occur in the Dutch Wadden Sea that can have detrimental effects on its inhabitants [85], such as commercial fishing, kite surfing, tourism, military exercises, and mining. In combination with large-scale phenomena, such as sea level rise and global warming [86], these anthropogenic activities can cause disturbances and habitat destruction, and thus contribute to population declines [87, 88]. The causes underlying the declines of shorebird population numbers in the Wadden Sea are often debated, partly because of our limited understanding of environmental processes and animal space use, which leads to tension and possibly conflict between stakeholders and management [61, 85, 89, 90]. The development of WATLAS has opened-up possibilities for quantifying space use of many small shorebirds directly, automatically, and at high spatiotemporal accuracy. WATLAS could thus aid in studies of impact assessment on shorebird space use, such as assessing the effects of disturbances and habitat loss due to land subsidence and sea level rise. More generally, WATLAS could facilitate evidence-based conservation, and aid the management of this UNESCO world heritage site.

## Conclusions

In this study, we introduced WATLAS as a high-utility tracking system in the Dutch Wadden Sea, capable of tracking hundreds of small individuals simultaneously at high spatiotemporal resolution. After the initial investment for an array of receivers (which can be substantial), the costs per tag are low, which facilitates regional, long-term studies on movement ecology and space use of many individuals and multiple species and facilitates collaboration between researchers across research institutes. Additionally, maintenance and operating costs need to be considered. Because tags are lightweight as well as cheap, WATLAS can facilitate studies on, for instance, collective behaviour, social information use, and movement ecology of entire communities of free-living animals. WATLAS can also support evidence-based nature conservation and management, for example with assessing the impact of anthropogenic activities on space use of shorebirds. More generally, with WATLAS, animals can function as sentinels informing us about the state of the Wadden Sea ecosystem [62, 76], and thus aid nature conservation and management of this globally important ecosystem.

## Supporting information

Additional file 1 with supplementary figures

Additional file 2 with animation of red kot movement

## DECLARATIONS

## Acknowledgements

Many people and organisations are involved in hosting the WATLAS equipment, without whom this study would be impossible. We therefore thank Hoogheemraadschap Hollands Noorderkwartier, Koninklijke Nederlandse Redding Maatschappij, Staatsbosbeheer, Marine Eco Analytics, Koninklijke Luchtmacht, Het Posthuys, Natuurmonumenten, Wetterskip Fryslan, Afsluitdijk Wadden Center, Vermilion, Rijkswaterstaat, Carl Zuhorn, Lenze Hofstee and Lydia de Loos. We thank Natuurmonumenten for access to Griend and using their facilities. Also, we thank Hein de Vries, Klaas Daalder, Hendrik-Jan Lokhorst, Bram Fey, Wim-Jan Boon from the RV Navicula and RV Stern, as well as the many other NIOZ staff and volunteers that facilitated this work. We would particularly like to thank Anita Koolhaas, Hinke and Cornelis Dekinga for their help with building the receiver stations. We thank Jeras de Jonge, Martin Laan, Sander Asjes, and Aris van der Vis for their technical help, and Benjamin Gnep for his beautiful photos and persistently replacing broken LNA’s. Thanks to Marten Tacoma for visualizing the tracking data in real-time on www.nioz.nl/watlas and Ingrid de Raad for help posting WATLAS-related news. We also thank the Minerva Foundation and the Minerva Center for Movement Ecology for supporting the development and maintenance of all ATLAS systems, and for Yotam Orchan and Yoav Bartan for their most valuable technical assistance. Finally, we thank Theunis Piersma, two anonymous referees and the associate editor for providing valuable comments and suggestions for improvements on an earlier version of this manuscript.

## Ethics approval and consent to participate

Permits to catch, handle and tag sanderlings and red knots was granted to the NIOZ under protocol number NIOAVD8020020171505.

## Consent for publication

Not applicable

## Availability of data and materials

Upon publication, the datasets supporting the conclusions of this article will be included within the article and its supplementary files.

## Authors’ contributions

AIB conceived the idea for the manuscript, analysed the data and led the writing of the manuscript; RN and ST developed the ATLAS system and provided support throughout data collection; FvM, BD and AD built and maintained the WATLAS system and tags, and provided operational support; AIB, AD, LdM, JtH, CEB and RAB deployed the receiver stations; AIB, AD, EP, SE, PG, LdM, JtH, RAB, and CEB caught and tagged birds; all authors contributed critically to the drafts and gave final approval for publication.

## Additional file information

Additional file 1

Microsoft Word file (.docx)

Supplementary figures S1-S5.

Additional file 2

MPEG animation (.mp4)

Animation of red knot movement near Griend between two consecutive high tides.

Different individuals are indicated with different colours. However, due to the large number of individuals (N=79), colours can be used more than once between individuals.

