## Additional file 1 with supplementary figures for "WATLAS: high throughput and real-time tracking of many small birds in the Dutch Wadden Sea"

^3^ Blavatnik School of Computer Science, Tel-Aviv University, Tel Aviv 67798, Israel

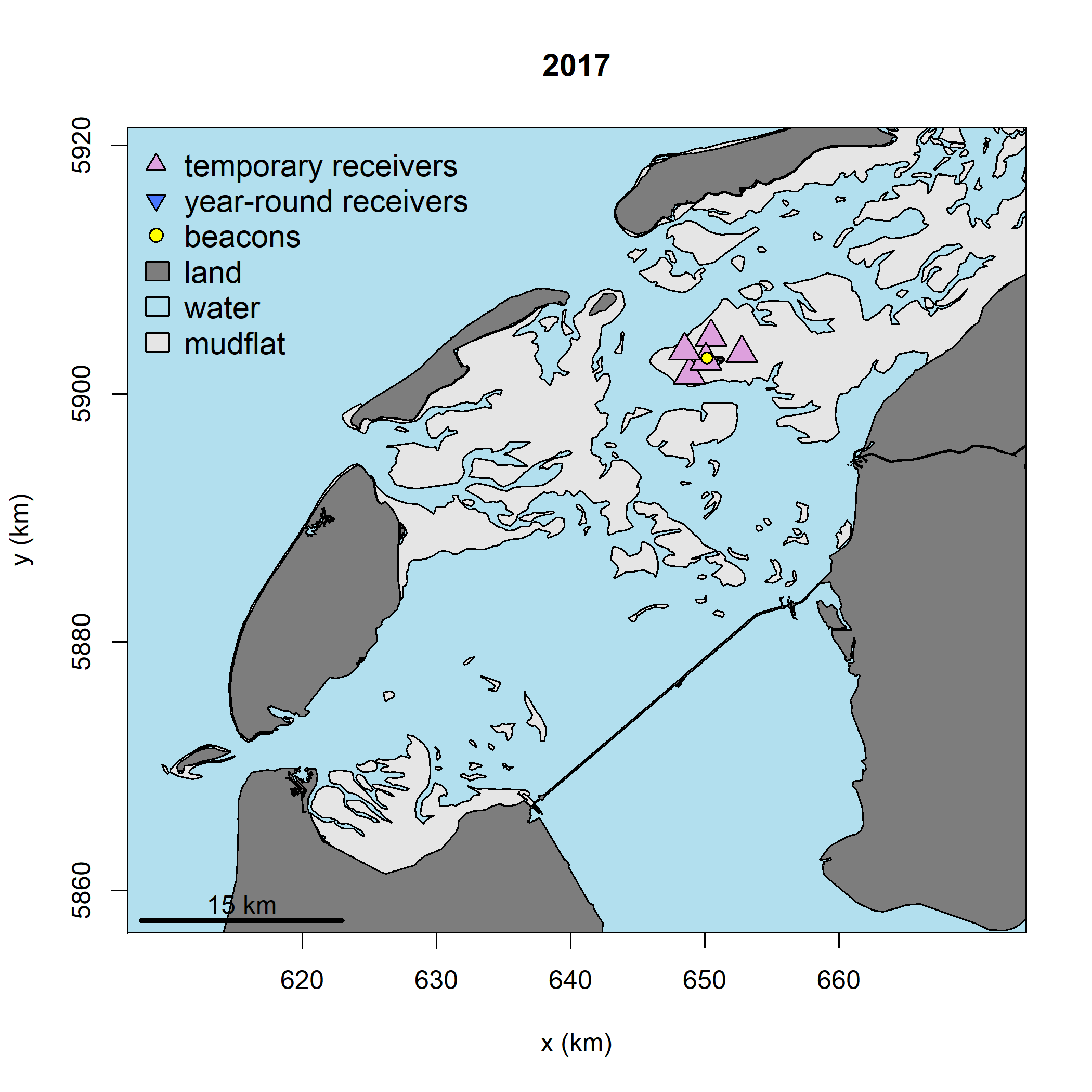

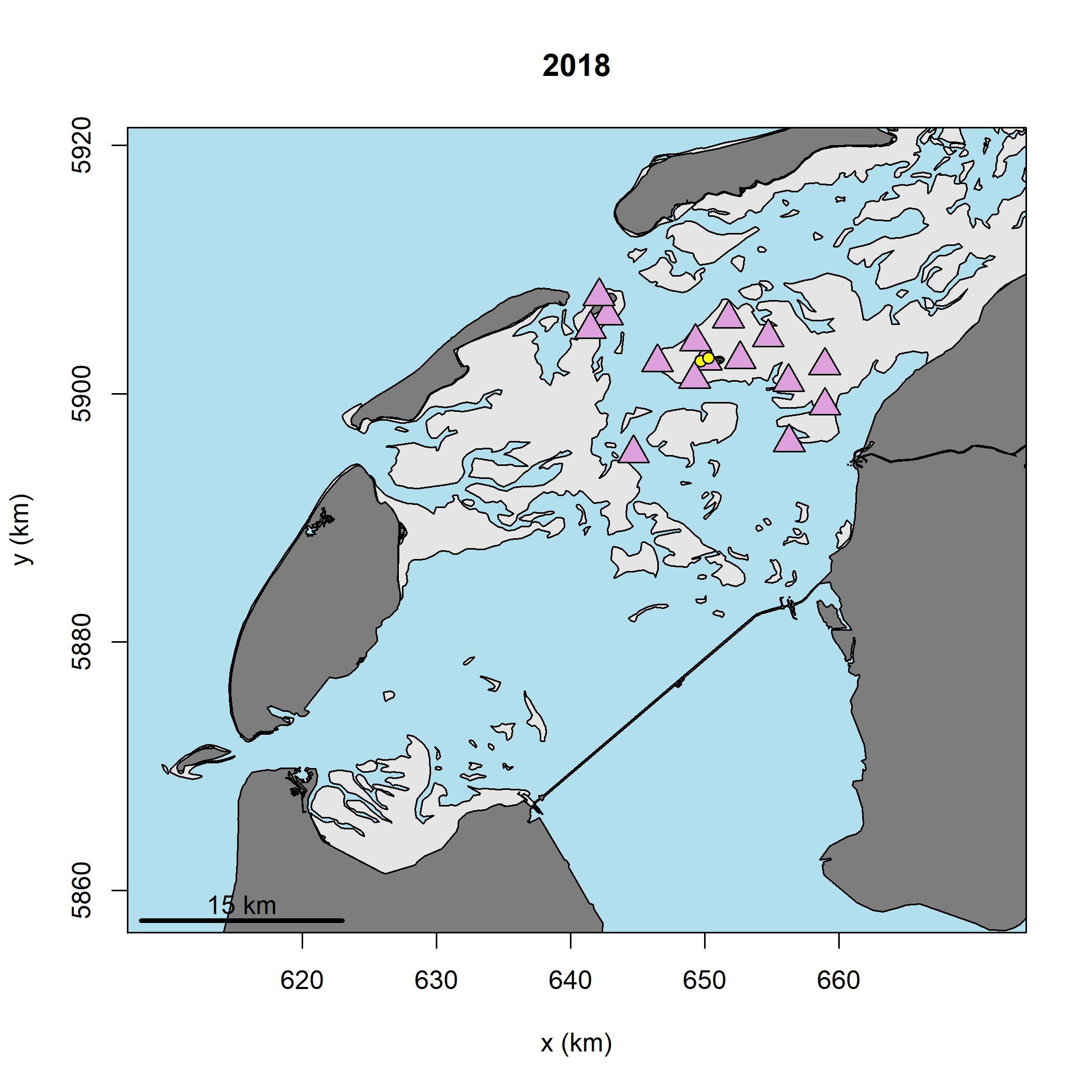

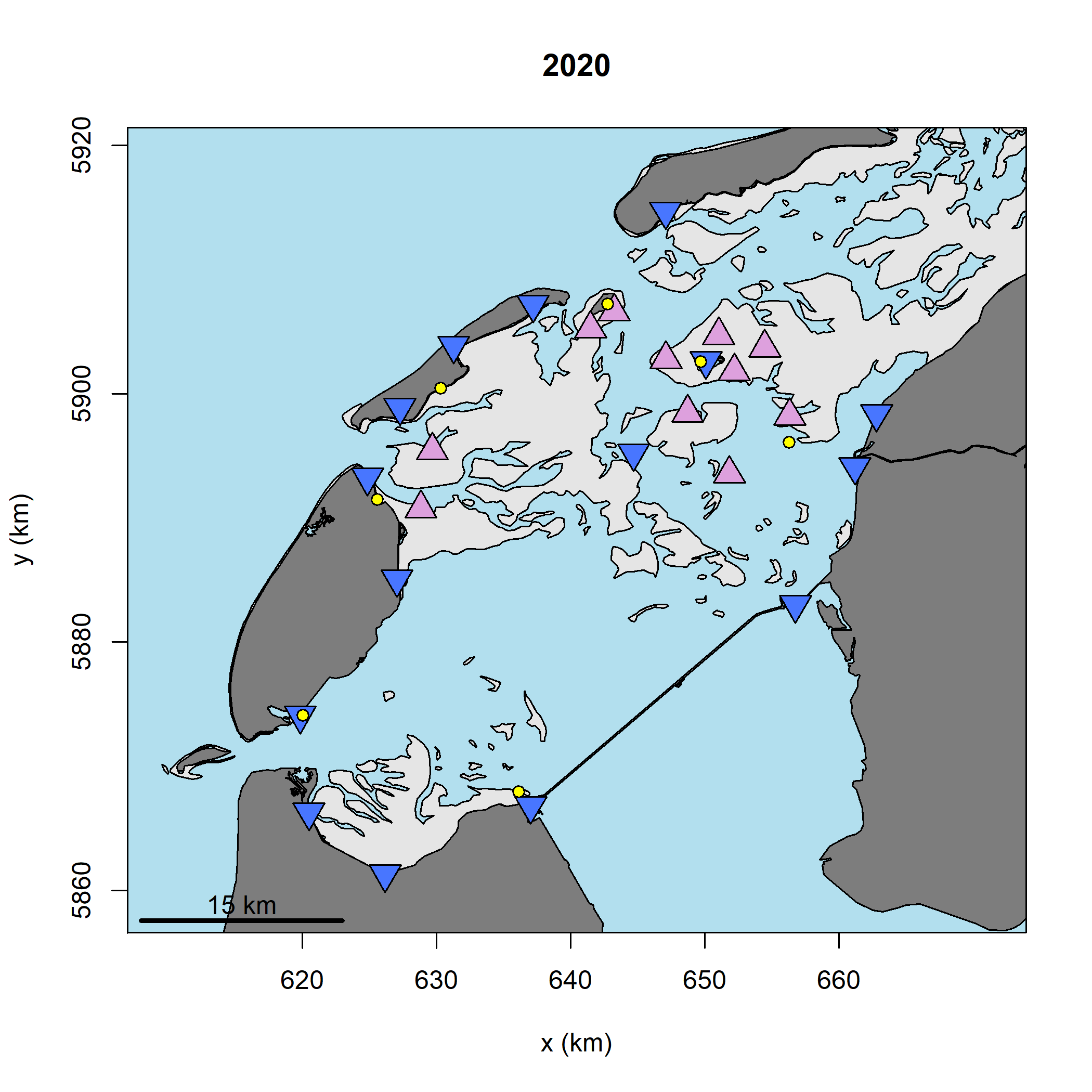

***Fig. S1*** *Maps of the receiver array and beacon locations in 2017, 2018 and 2020. Land is shown in dark grey, water in blue and mudflat in light grey. See Fig. 1 for the receiver array in 2019. The coordinate system refers to UTM 31N. © map data from Rijkswaterstaat.*

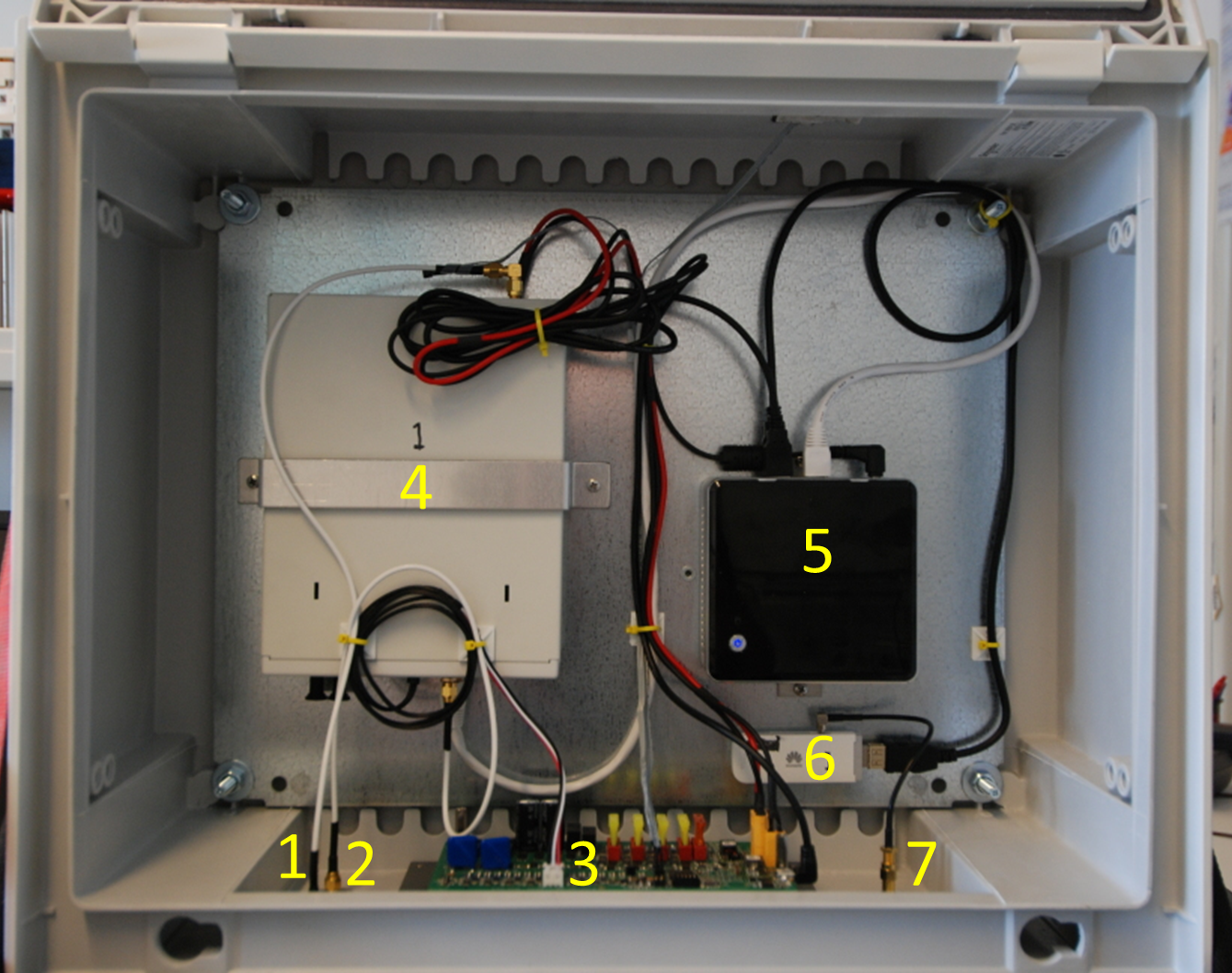

***Fig. S2*** *The inside of a receiver cabinet: 1) incoming connection from external GPS antenna used to synchronize clocks, 2) input from external main antenna, 3) circuit board that controls the power from solar panels and wind generators, 4) software defined radio, 5) on board computer, 6) mobile dongle with SIM-card, and 7) antenna for 3G communication.*

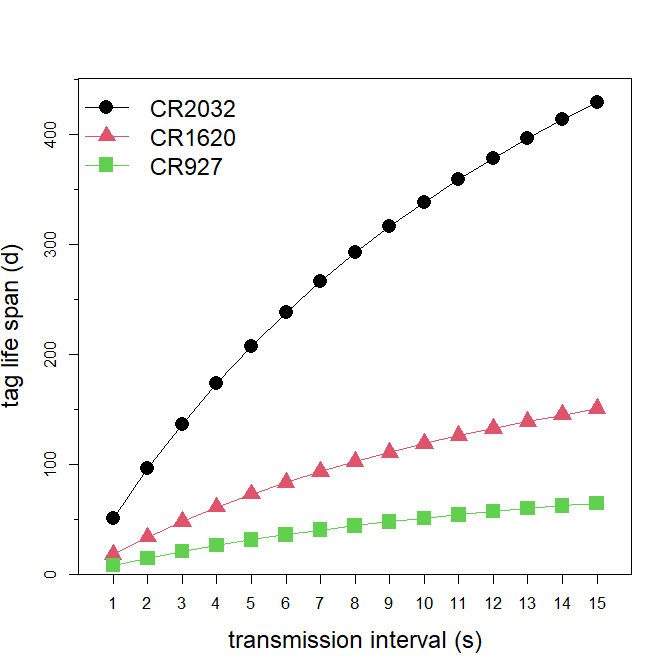
***Fig. S3*** *Estimated tag life- span at different transmission intervals for battery types CR2032, CR1620 and CR927 that respectively weigh 3.3, 2.0 and 0.6 g. To calculate the life span of tags, the manufacturer-specified capacity of the batteries (respectively 30, 70, and 220 mAh) was reduced by 10 % because their capacity will likely not be fully utilized.*

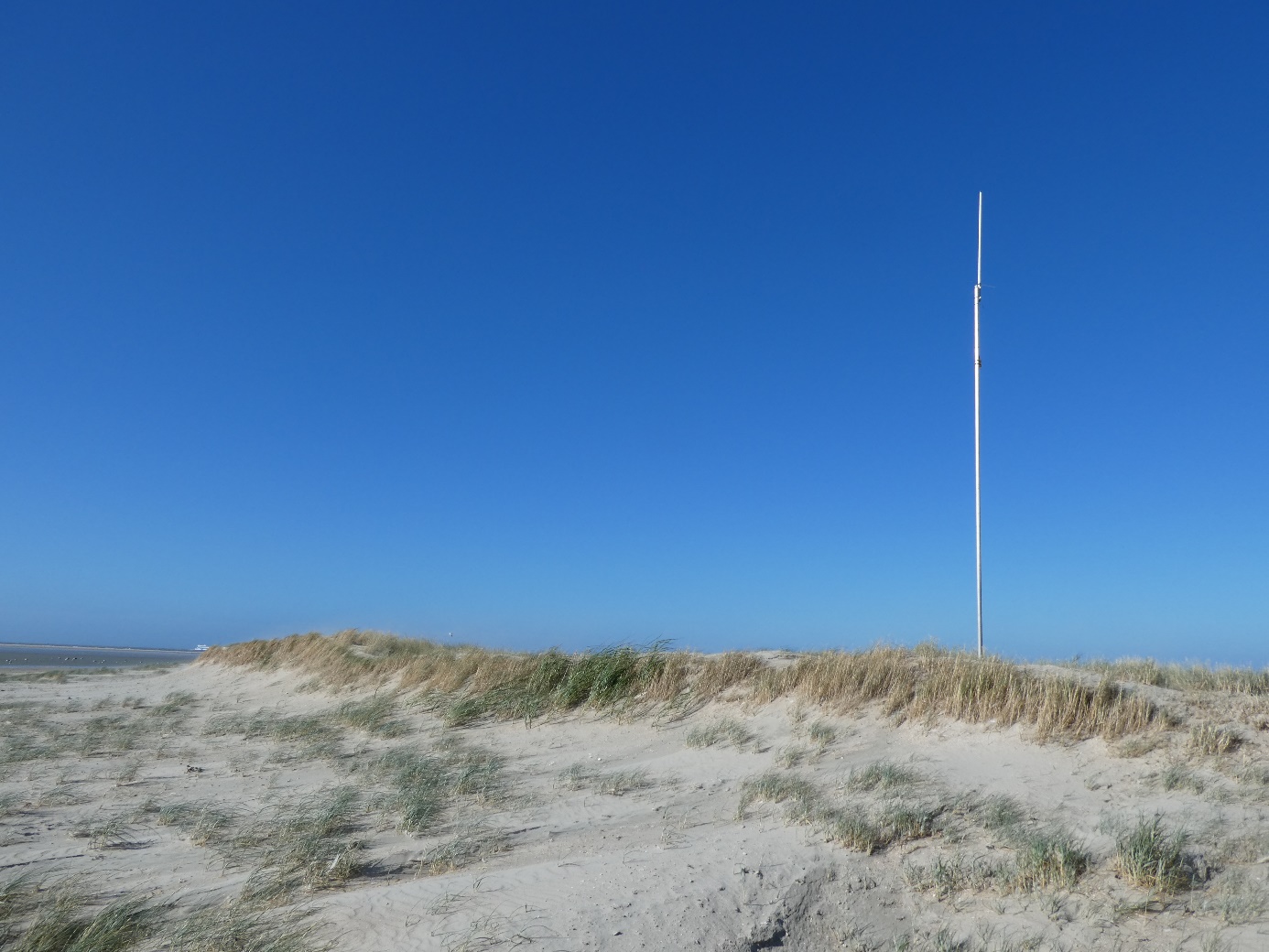

***Fig. S4*** *One of seven beacons mounted on a 6 m aluminium scaffold.*

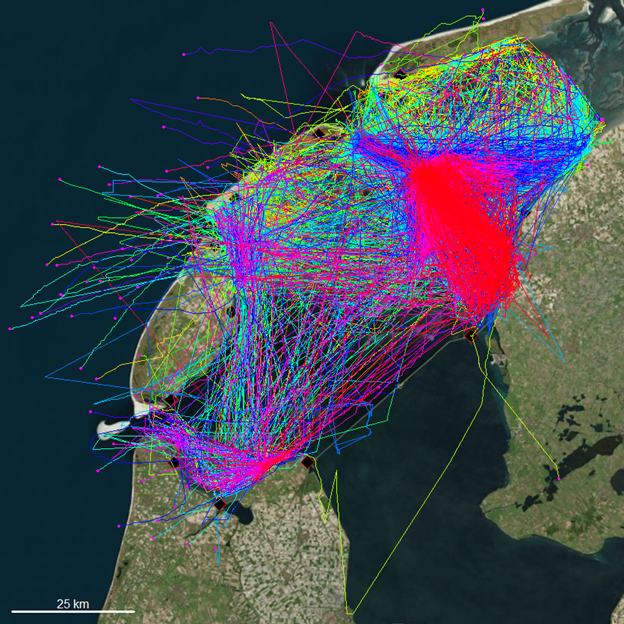

***Fig. S5*** *The tracks of 221 red knots tracked between between 1 August and 1 November 2019, coloured by individual. The magenta coloured points indicate the final position of a bird.*
